# Serine hydroxymethyl transferase is a binding target of caprylic acid: Uncovering a novel molecular target for a herbicide and for producing caprylic acid-tolerant crops

**DOI:** 10.1101/2023.12.12.571245

**Authors:** Zuren Li, Mugui Wang, Haodong Bai, Hongzhi Wang, Jincai Han, Likun An, Dingfeng Luo, Yingying Wang, Wei Kuang, Xiaoyi Nie, Lianyang Bai

## Abstract

Identification of new binding targets is essential for the development of herbicides and phytotoxin-tolerant crops. Caprylic acid (CAP) is a safe and non-selective bio-herbicide in uncultivated areas. However, the herbicidal action of CAP remains unclear. Herein, metabolomic and proteomic profiling indicated that a serine hydroxymethyl transferase in *Conyza canadensis* (*Cc*SHMT1) is a promising candidate binding targeted for CAP. The protein abundance and activity of *Cc*SHMT1 were decreased in a time- and dosage-dependent manners after CAP treatment. CAP competes with phenyl-serine at the binding sites, decreasing the enzymatic activity of *Cc*SHMT1. Overexpression of *CcSHMT1, AtSHMT1* and *OsSHMT1* in *Arabidopsis* or rice endowed plants with high tolerance to CAP treatment, whereas the knockout of *osshmt1* led to death of plants under normal atmospheric conditions. Furthermore, T194A, T194A/ M195V and D209N *Osshmt1* mutant plants derived from base editing exhibited tolerance to CAP. CAP bound to *Cc*SHMT1 with high affinity, and Ala191 in the key domains of N-terminus was identified to be critical for the binding site of CAP. Collectively, our findings demonstrate that *Cc*SHMT1 is a binding target for herbicidal activity of CAP. This study marks a key step in the druggability of SHMT inhibitors and represents an attractive target for phytotoxin-tolerant crops.

## Introduction

Herbicide application is the most effective and affordable weed management strategy (Gerhards et al. 2011). However, of the 25 known herbicide targets, weeds resistant to 21 targets have been reported, including 267 weed species that often appear in 96 crops (Foucart and Horel 2018; Heap 2023). To overcome this challenge, application of new herbicides with novel modes of action is a good strategy to overcome or delay herbicide resistance. However, very few novel herbicidal targets have been discovered in recent decades (Duke and Dayan 2022).

Phytotoxins have emerged as novel herbicide candidates targeting distinct plant processes (Macias et al. 2019). Several medium-chain fatty acids, such as pelargonic acid (CH₃(CH₂)₇CO₂H), were identified as effective, broad-spectrum phytotoxins with low risk for development of herbicide resistance (Coleman and Penner 2008; Real et al. 2021). The mechanisms of action of phytotoxins can be divided into non-specific and specific modes of action, similar to other abiotic stresses (salinity or alkalinity stress) (Zhu 2016). We recently investigated caprylic acid (C_8_H_16_O_2_, CAP), which exhibits high herbicidal activity by destroying chloroplasts and mitochondria in leaf cells and is a potent herbicide against weeds (Li et al. 2018; Li et al. 2019b). High doses of CAP typically destroy plant plasma membranes via lipid peroxidation, increasing singlet oxygen production and peroxidation reactions and causing necrosis (Fukuda et al. 2004; Lederer et al. 2004). These study presented a non-specific mode of action. However, the initial target enzymes of CAP remain unclear. The herbicidal binding targets of CAP maybe novel for uncovering a specific mode of action of phytotoxicity.

A combination of modern molecular, structural, and genomic technologies is considered as one of the best approaches for identifying herbicide targets (Duke et al. 2019). Structural analyses have shown that *Arabidopsis thaliana,* acetohydroxyacid synthase (AHAS) form tight complexes with pyrimidinyl-benzoate (PYB) and sulfonylamino-carbonyl-triazolinone (SCT) herbicides, providing a structural basis for designing new AHAS-targeting herbicides (Garcia et al. 2017; Lonhienne et al. 2020). High-throughput screening of target enzymes has been employed to identify novel herbicides such as aspterric acid (Lein et al. 2004; Yan et al. 2018). Proteomics has been used to confirm low enzyme-level molecular sites of natural herbicidal compounds (Dayan et al. 2020). However, using these approaches to identify novel herbicidal targets is still scarce.

In a previous study, we identified 112 differentially expressed proteins in CAP-treated *Conyza canadensis*, including G0WXY7, A0A103TF61 and A0A124SCM3 (Li et al. 2019a). However, the exact binding targets of CAP were not discerned. In the present study, we used untargeted metabolomics to identify different metabolites after CAP treatment, including glycine, palmitic acid and vanillin. We then co-analyzed the proteomic and metabolomic data to narrow down the candidate CAP targets, including SHMT1, FBA5, PGK3, PGK2, PGM1, GLO1, CYFBP, ADK2, and PED1. These analyses identified a serine hydroxymethyl transferase of *C. canadensis* (*Cc*SHMT1) as a potential herbicide target of CAP. To validate this, we analyzed *Arabidopsis* and rice lines overexpressing heterologous *CcSHMT1* with CAP. Subsequently, the coding sequences of *OsSHMT1* were *in vivo* edited to obtain tolerant crops using base editing. The homology modeling structure of *Cc*SHMT1, which shows 87.97% sequence identity with that of *At*SHMT2 (PDB: 6SMN.1.A), was used for *in silico* docking of CAP to identify the amino acid residues that are critical for binding. The present study uncovered a CAP-binding target enzyme and provided an integrated theoretical framework for herbicidal target discovery.

## RESULTS

### *CcSHMT1* is a candidate action target of CAP

The phytotoxic symptoms appeared after 4 h treatment at the 625 μM CAP (Li et al. 2019c). To identify the targets of CAP, we first employed untargeted metabolomics to analyze the significantly different metabolites at 4 h post-treatment by 625 μM CAP from the untreatment. Principal component analysis (PCA) and multivariate score plots of all the samples and quality controls (QCs) showed a reliable trend of intergroup separation (R_2_X= 0.577> 0.5), good predictive power, and goodness-of-fit (Supplemental Table 1S and Fig. 1S). Differentially expressed metabolites (DEMs) were defined using two major parameters: a variable influence on projection (VIP) >1 and Student’s t-test *p*-value < 0.05. CAP treatment resulted in 151 DEMs (100 upregulated, 51 downregulated) at 4 h post-treatment relative to the untreated controls (Fig. 1A, Supplemental Table 2S). The DEMs were enriched in various amino acid metabolic pathways, including carbon fixation in photosynthetic organisms, carbon metabolism, and glyoxylate and dicarboxylate metabolism (Fig. 1B). The amino acid metabolic pathways of Val, Leu, and Ile and Gly, Ser, and Thr biosynthesis are closely associated with glyoxylate and dicarboxylate metabolism pathways. Our previous proteomic analysis showed that the abundance of 112 proteins either increased or decreased after CAP treatment, and among these proteins, among which G0WXY7 (Ribulose bisphosphate carboxylase large chain, involved in carbohydrate metabolism), A0A103TF61 (Aspartic peptidase, involved in hydrolase activity) and A0A124SCM3 (Electron transport accessory protein-like domain-containing protein, involved in photosystem I) exhibited the most pronounced changes (Li et al. 2019a).

**Fig. 1.**
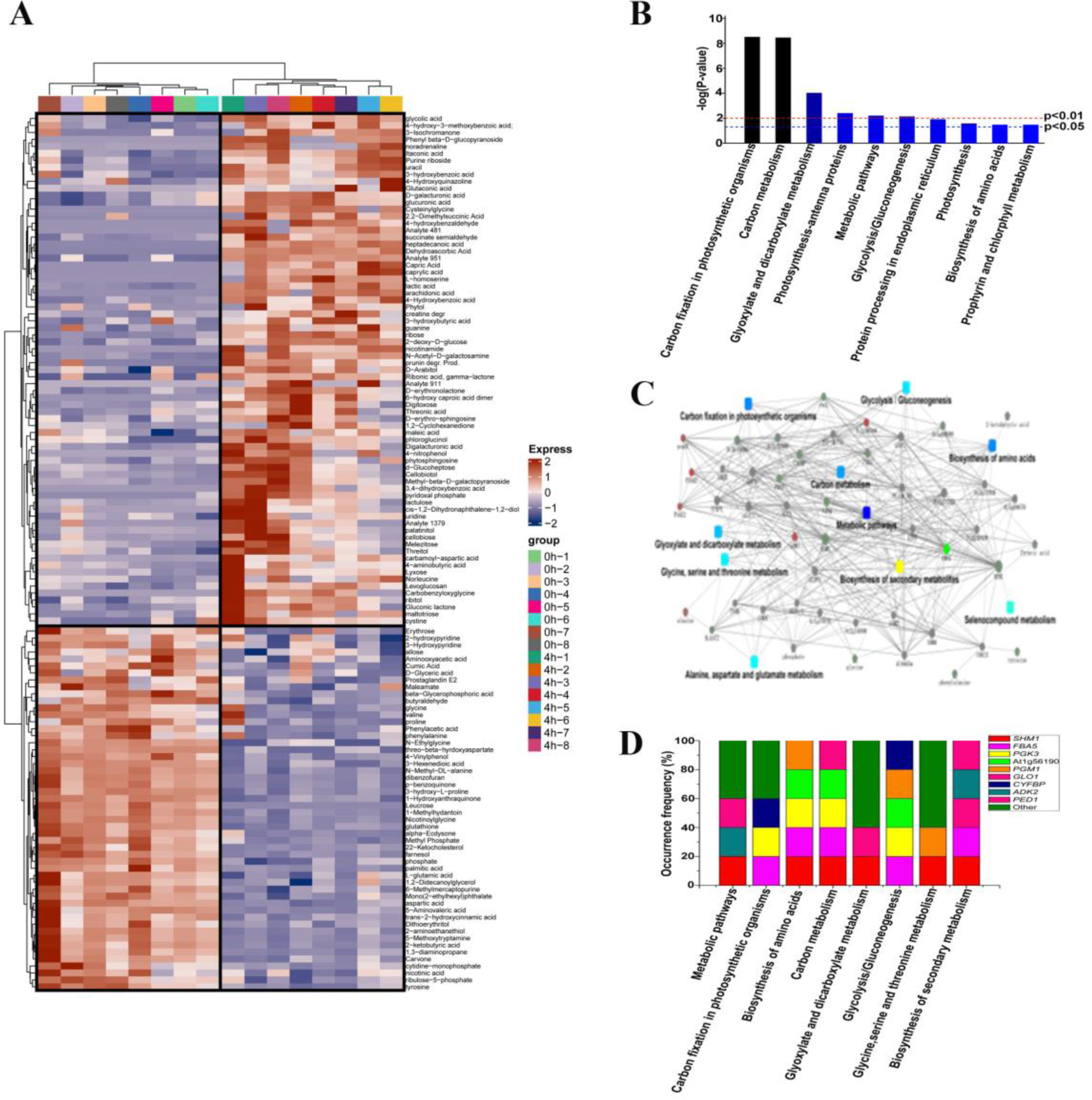
Metabolomic profiling of C. canadensis at various timepoints post caprylic acid (CAP) treatment versus the untreated control. At 4 h after foliar application of CAP treatment (625 μM) and untreated control, a total of 2 g of *C. canadensis* leaves were collected for analysis by GC-MS. (A) The heatmap analysis of significantly different metabolites The complete list of perturbed metabolites is in Supplemental Table 2S, Student’s t-test *p* valure<0.05; (B) The top ten pathways affected KEGG Metabolic pathway by CAP treatment after proteomic and metabolomic co-analysis. C: metabolomics and proteomics data correlation pathway analysis. (D)The occurrence frequencies of the top nine genes involved in the top eight KEGG pathways were key indicators of the differences between CAP-treated samples and untreated controls. The occurrence frequencies of the genes increased by 20% and appeared at least once in the KEGG pathways, according to the standard calculating module.

The availability of both proteomic data (Li et al. 2019a) and metabolomic data prompted us to perform a comparative KEGG pathway analysis. KEGG pathway enrichment analysis showed that the proteins and metabolites that changed after CAP treatment were mainly enriched in 10 pathways, including carbon fixation in photosynthetic organisms, amino acid biosynthesis, and carbon metabolism (*P*< 0.05) (Fig. 1C). The expression of nine genes (*SHMT1*, *FBA5*, *PGK3*, *PGK2*, *PGM1*, *GLO1*, *CYFBP*, *ADK2*, and *PED1*), which are known to participate in the top eight pathways, was significantly different between CAP-treated samples and untreated controls. The occurrence frequency of all these genes that appeared at least once in the top eight KEGG enriched pathways was increased by 20% according to the standard calculating module. The standard calculation module is similar to the occurrence frequency of herbicide resistance as an indicator of harmful and resistant weeds (Walsh et al 2001).

Notably, a serine hydroxymethyltransferase 1 (*CcSHMT1*) showed the highest frequency among the nine genes and was involved in six of the eight pathways in *C. canadensis* (Fig. 1D). SHMT1 plays a role in photorespiration and metabolic conversion of glycine to serine (Douce et al 2001; Bauwe and Kolukisaoglu 2003). Specially, two amino acids were significantly decreased after CAP treatment (Supplemental Fig. 2S). Hence, multiple omics analyses suggested that *Cc*SHMT1 is the most likely binding target of CAP.

### CAP inhibited SHMT activity but upregulated *CcSHMT1* expression

Subsequently, we assessed transcriptional alterations in *SHMT* genes and SHMT enzyme activity in *C. canadensis* to corroborate the omics results. We first characterized one of the *C. canadensis SHMT* gene, *CcSHMT1* (Fig.2A), which is highly similar to its orthologs in other plant species (Fig.2B) (Lakhssassi et al. 2019). The protein-coding region of *CcSHMT1* was 1,542 bp in length (MN586851.1), encoding a protein of 513 amino acids with a molecular weight of 56.994 kDa and a theoretical pI of 8.82. *Cc*SHMT1 was located in the mitochondria, with GFP fluorescence overlapping with Mito fluorescence, which was identical to that reported for SHMT1(Fig. 2C) (Wang et al. 2015; Spalding et al. 2010; Zhou et al. 2015). In addition, we resolved the homology model structure of *Cc*SHMT1 in AlphaFold2 (Baek et al. 2021). The homologous structure showed 88.03% sequence identity with *Arabidopsis* SHMT2 structure (PDB: 6SMN). The overall three-dimensional (3D) homologous structure of *Cc*SHMT1 forms a tetramer with four identical subunits. The monomer contained three domains: N-terminal, large domain, and C-terminal (Fig. 2D). Hence, *Cc*SHMT1 is a typical PLP domain-containing enzyme.

**Fig. 2.**
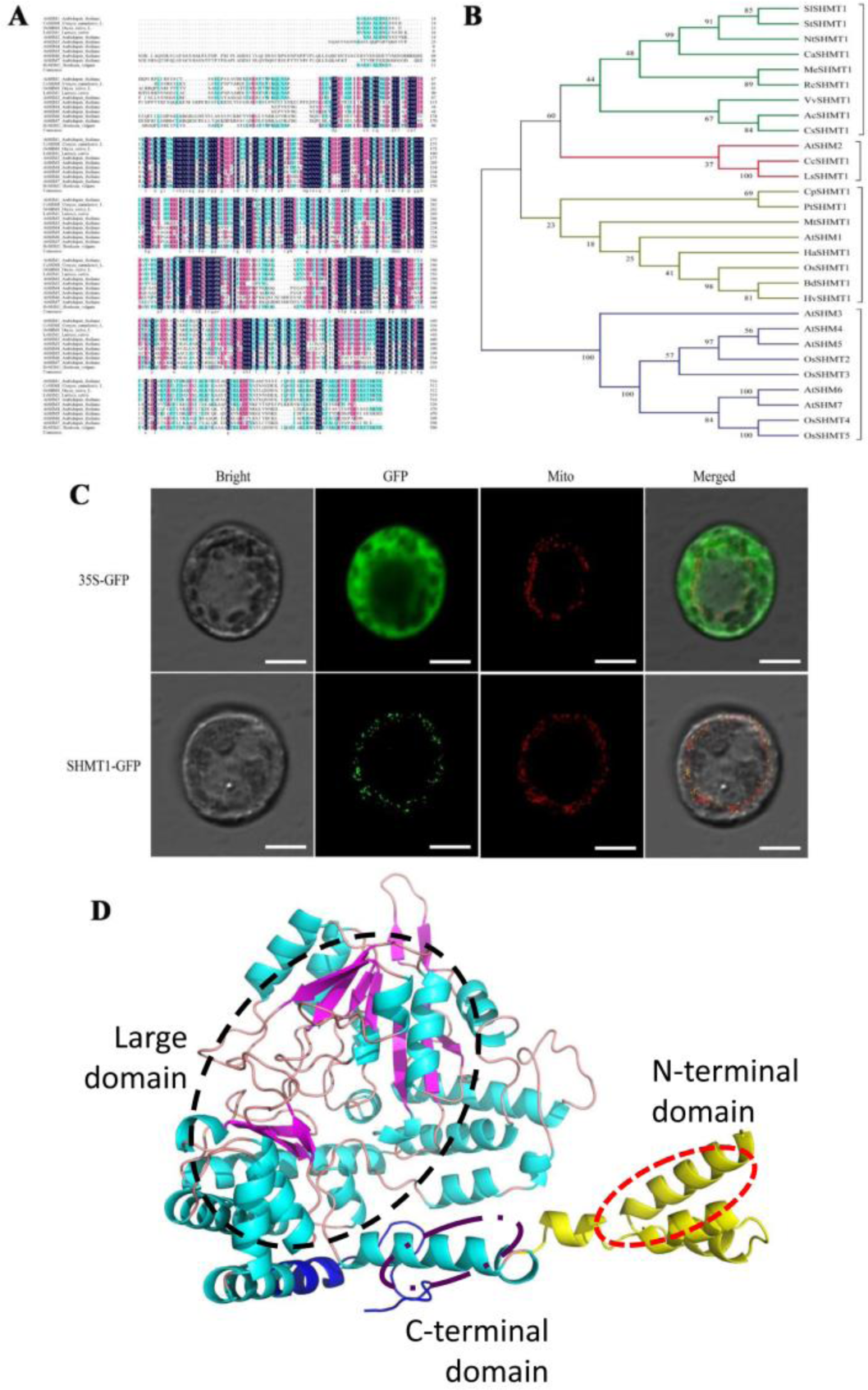
*Cc*SHMT1 is a typical PLP enzyme. (A) Multiple sequence alignment of SHMTs. The sequences in the boxes are conserved amino acid residues. Source species of SHMT proteins and their GenBank accession numbers: BdSHM1, *Brachypodium distachyon*, XP_003559966.1; CcSHM1, *Conyza canadensis*, MN586851.1; HaSHM1, *Helianthus annuus*, XM_022184227.1; NtSHM1, *Nicotiana tabacum*, XP_016446471.1; HvSHM1, *Hordeum vulgare*, KAE8805790.1; AtSHM1, *Arabidopsis thaliana,* NC_003075.7; OsSHM1, *Oryza sativa*. (B) A molecular phylogenetic tree of SHMT gene based on amino acid sequences with homologous proteins from other species (neighbor-joining method, 1, 000 replicates). The sequences were obtained from GenBank. Source species of SHMT proteins and their GenBank accession numbers. (C) The subcellular Localization of *CcSHMT1*, the bar*=*10 μm. (D) The homology modeling structure of *Cc*SHMT1.

A quantitative real-time polymerase chain reaction (qRT-PCR) assay showed that the transcription level of *CcSHMT1* was elevated more than five-fold within 30 min after CAP treatment. A few other genes were also upregulated to some degree, including *FBA5*, *PGK3*, *PGM1,* and a homolog of *At1g56190*, but to much smaller magnitudes (Fig. 3A). Additionally, *CcSHMT1* was only upregulated for a short time, as its transcription level peaked at 1 h after CAP treatment and then sharply decreased to a steady-state level 2 h after treatment (Fig. 3B). We also measured the protein level of soluble SHMT in *C. canadensis* using a commercially available SHMT ELISA kit (Mei5 Biotech., Beijing, China). The total SHMT protein level increased gradually from 0.5 h to 2 h post-treatment, reaching its highest level (364.53 U/L) at 2 h, and then began to decrease (Fig. 3C). These results suggested that induction of *CcSHMT1* was a transient response to CAP treatment.

**Fig. 3.**
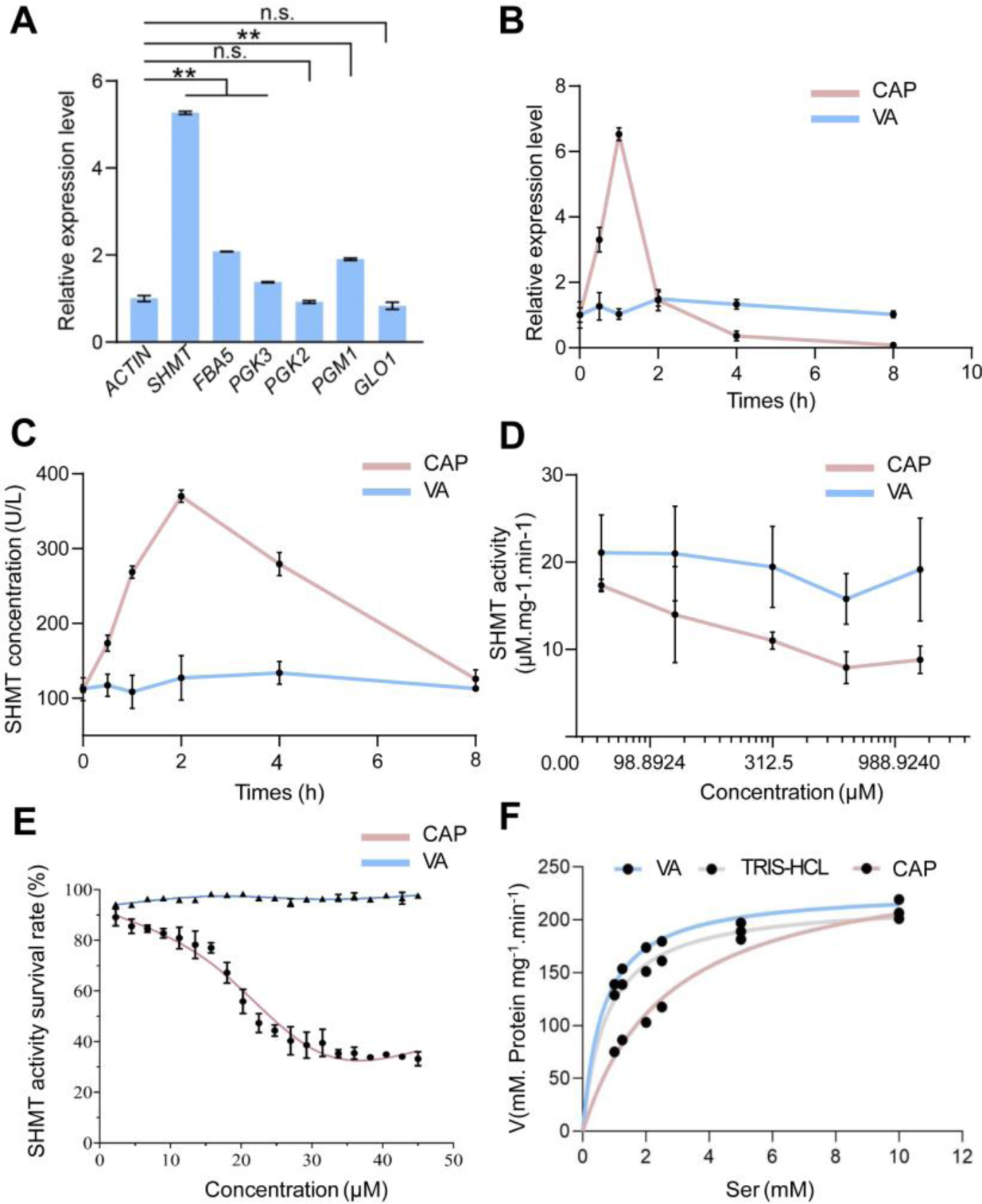
*CcSHMT1* gene expression, protein abundance, and enzyme assays of CAP treatment versus the untreated control. (A) Relative expression of the top five genes in *C. canadensis* following CAP treatment. Primer-Q were designed by DOMAN 7 according to the conserved sequences from NBCI BLAST; *SHMT1*, *FBA5*, *PGK3*, *PGK2*, *PGM1* and *GLO1* were had homologous genes from *Arabidopsis*; the mRNA was extracted from *C. canadensis* 30 min after CAP treatment (625 μM). (B) Relative expression of *CcSHMT1* in *C. canadensis* under CAP and Vanillic acid (VA) treatments. The mRNA was extracted from *C. canadensis* 0, 0.5, 1, 2, 4, and 8 h after CAP and VA treatment (625 μM). (C) Effect of *Cc*SHMT1 concentration *in vivo* under CAP and VA stress. The *Cc*SHMT1 protein was extracted from *C. canadensis* leaves 0, 0.5, 1, 2, 4, and 8 h after CAP and VA treatment (625 μM). The protein levels were measured with a SHMT ELISA Kit according to 5 the manufacturer’s instructions. (D) Effect of *Cc*SHMT1 activity *in vivo* under CAP and VA stress. The *Cc*SHMT1 protein was extracted from the *C. canadensis* leaves after CAP and VA treatment (0, 78.125, 156.25, 312.5, 625, 1250 μM). The SHMT activity was assayed based on the product rate of benzaldehyde. (E) The effect of *Cc*SHMT1 activity *in vitro* under CAP and VA stress. Recombinant *Cc*SHMT1 was purified using Ni^2+^-chelating columns. SHMT activity was assayed by the product rate of benzaldehyde 0.5 h after CAP and VA treatment (0.63, 0.95, 1.26, 1.58, 1.90, 2.53, 3.16, 6.32, 9.48, 12.64, 15.80, 18.96, 22.12, 25.28, 28.44, 31.60, 34.76, 37.92, 41.08, 44.24 μM). (F) Kinetic analysis of *Cc*SHMT1 *in vitro* under CAP and VA stress. Recombinant *Cc*SHMT1 proteins were purified with Ni^2+^-chelating columns, and kinetic analyses for DL-β-phenylserine were performed (1 mM, 1.25 mM, 2.00 mM, 2.5 mM, 5 mM, 10 mM of DL-β-phenylserine). CAP: caprylic acid, VA: Vanillic acid. Each independent measurement was repeated at least three times. Data are the mean±standard deviation (s.d.). Statistically significance was at *p*-value < 0.05 according to the Student’s t-test, *P<0.05, **P<0.01, ***P<0.001 and NS, not significant (P>0.05).

Using the same *C. canadensis* samples, the total SHMT activity was examined based on the production rate of benzaldehyde. SHMT activity decreased gradually to 17.331 μmol phenylserine·min^-1^·mg^-1^ protein after treatment of 78.125 μM CAP, compared with untreated samples (19.901 μmol phenylserine·min^-1^·mg^-1^ protein) and Vanillic acid (VA, C_8_H_8_O_4_) treated samples (21.080 μmol phenylserine·min^-1^·mg^-1^ protein), and CAP inhibited SHMT activity in a dose-dependent manner (from 78.125 μM to 1250 μM) (Fig. 3D). However, the activity of the photorespiratory GLO1, the transcript levels of which showed similar changes as SHMT1 under CAP treatment *in vivo*, was not significantly inhibited by CAP (Supplemental Fig. 3S). Taken together, these data indicated that CAP treatment induced transcription of *CcSHMT1*, which was positively correlated with higher protein level during a brief post-treatment time window.

To determine whether CAP could inhibit the enzymatic activity of SHMT *in vitro*, *Cc*SHMT1 was expressed in *Escherichia coli* and purified using Ni^2+^-chelating columns (Supplemental Fig.4S). The activity of the purified *Cc*SHMT1 was analyzed by measuring the production rate of benzaldehyde in the absence or presence of CAP (Waditee-Sirisattha R et al. 2012). SHMT activity gradually decreased along with increase of CAP concentration from 0.63 to 44.23 μM, indicating that CAP dramatically inhibited SHMT activity in a concentration-dependent manner (Fig. 3E). The half-maximal inhibitory concentration (IC_50_) was 23.1 μM. In contrast, the fatty acid-controlled, VA did not suppress SHMT activity under the same conditions.

**Fig. 4.**
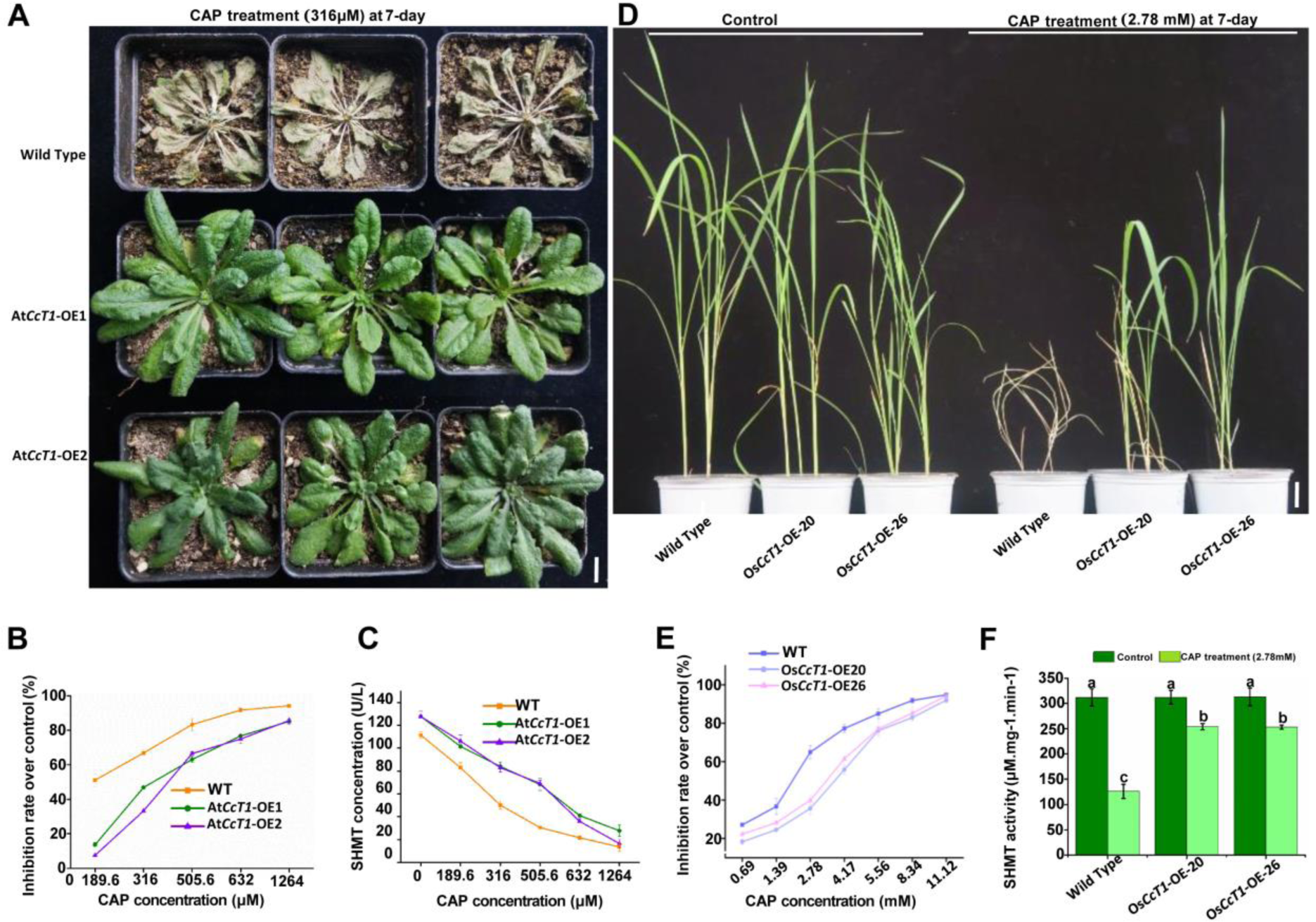
The *CcSHMT1*-overexpressing and WT in *Arabidopsis* and rice under CAP treatment versus the untreated control. (A) The two lines of *CcSHMT1*-overexpressing *Arabidopsis* plants (At*Cc*T1OE1 and At*Cc*T1OE2) and WT 7 days after 316 μM CAP treatment. (B) The inhibition rate of *CcSHMT1*-overexpressing and WT seedlings treated with CAP (189.6, 316, 505.6, 632 and 1264 μM). The inhibition rate was calculated after 7 days relative to the untreated control plants (Inhibition rate % = (Fresh Weight of untreated control plants-Fresh Weight of treatment plants)/ Fresh Weight of untreated control plants * 100%). The bioactivity assays comprised three biological replicates, each with three technical replicates. (C) The SHMT ac in *CcSHMT1*-overexpressing and WT *Arabidopsis* seedlings under CAP treatment (189.6, 316, 505.6, 632 and 1264 μM). (D) The two lines of *CcSHMT1*-overexpressing *Nipponbare* plants (Os*Cc*T1OE20 and Os*Cc*T1OE26) and WT 7 days after 2.78 mM CAP treatment. (E) The inhibition rate of *CcSHMT1*-overexpressing and WT seedlings treated with CAP (0.69, 1.39, 2.78, 4.17, 8.34 and 11.12 mM). The inhibition rate was calculated after 3 days relative to the untreated control plants. Data are the mean±standard deviation (s.d.). Statistically significance was at *p*-value < 0.05 according to Student’s t-test, *P<0.05, **P<0.01, ***P<0.001 and NS, not significant (P>0.05).

The kinetics properties of *Cc*SHMT activity under CAP and VA stress was comparatively its *V_max_* for DL-β-phenylserine under CAP treatment (238.10 μmol phenylserine·min^-1^·mg^-1^ protein) was not significantly different from that under VA treatment (227.27 μmol phenylserine·min^-1^·mg^-1^ protein) (Fig.3F, Supplemental Fig.5S). However, the *K_m_* of *Cc*SHMT1 under CAP treatment (2.24 mM) was higher than that under VA treatment (0.61 mM). Moreover, the *V_max_* and *K_m_* of untreat *Cc*SHMT1 was 212.76 μmol phenylserine·min^-1^·mg^-1^ protein and 0.68 mM. The *Km* value decrease with an invariable *Vmax* value resulted from CAP indicate that CAP is a competitive inhibitor of *Cc*SHMT1.

**Fig. 5.**
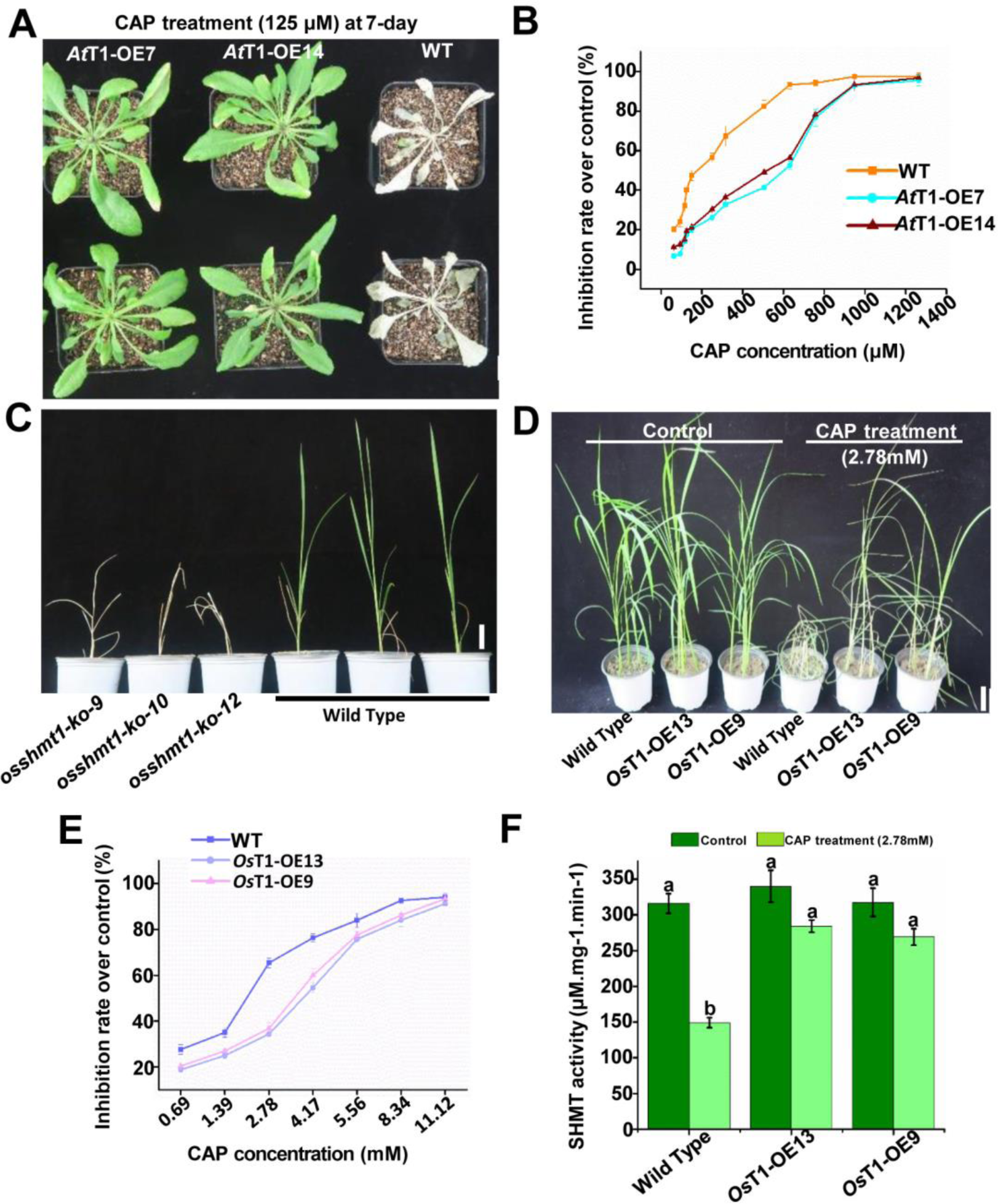
The *SHMT1*-overexpressing *Arabidopsis* and rice under CAP treatment versus the untreated control. (A) The two lines of *AtSHMT1*-overexpressing *Arabidopsis* plants (*At*T1-OE1 and *At*T1-OE2) and WT 7 days after 150 μM CAP treatment. (B) The inhibition rate of *AtSHMT1*-overexpressing and WT seedlings treated with CAP (62.5, 93.75, 115.62, 125, 150, 250, 316, 505.6, 632, and 1264 μM). The inhibition rate was calculated after 7 days relative to the untreated control plants (Inhibition rate % = (Fresh Weight of untreated control plants-Fresh Weight of treatment plants)/Fresh Weight of untreated control plants * 100%). The bioactivity assays comprised three biological replicates, each with three technical replicates. (C) The typical phenotype *OsSHMT1* knock-out mutant lines (*osshmt1-qc-9*, *osshmt1-qc-10*, *osshmt1-qc-12*). Bar= 2cm (D) The two lines of *OsSHMT1*-overexpressing *Nipponbare* plants (*Os*T1-OE13 and *Os*T1-OE9) and WT 7 days after 2.78 mM CAP treatment. Bar= 5cm (E) The inhibition rate of *CcSHMT1*-overexpressing and WT seedlings treated with CAP (0.69, 1.39, 2.78, 4.17, 8.34 and 11.12 mM). The inhibition rate was calculated after 3 days relative to the untreated control plants. (F) The SHMT activity of *CcSHMT1*-overexpressing and WT seedlings treated with CAP (2.78 mM). Data are the mean±standard deviation (s.d.). Statistically significance was at *p*-value < 0.05 according to Student’s t-test, *P<0.05, **P<0.01, ***P<0.001 and NS, not significant (P>0.05).

### Overexpression of *CcSHMT1* in *Arabidopsis* and rice increased resistance to CAP

To verify whether CAP targets *Cc*SHMT1, we transformed *CcSHMT1* into *Arabidopsis* and rice plants. Transgenic *Arabidopsis* (At*CcSHMT1*-OE) and rice (Os*CcSHMT1-*OE) were confirmed using PCR (Supplemental Fig.6S). RT-qPCR assays showed that the transcription level of *SHTM1* was highly increased in the At*CcSHMT1*-OE and Os*CcSHMT1-*OE lines (Supplemental Fig. 7S). To optimize the concentration of CAP for the treatment of *Arabidopsis*, At*CcSHMT1*-OE, Os*CcSHMT1-*OE, and wild type (WT) seedlings were treated with CAP at different concentrations. Upon treatment with CAP at 316 μM, two At*CcSHMT1*-OE lines did not show visible injury, while the WT plants were severely injured and did not survive beyond 7 days post-treatment (Fig. 4A). For the two independent At*CcSHMT1*-OE lines, the half-maximal effective concentration (EC_50_) of CAP was 376.3 μM (At*CcSHMT1*-OE1) and 413.9 μM (At*CcSHMT1*-OE2), whereas the EC_50_ for WT plants was 190.8 μM (Fig. 4B, Supplemental Table 3S). Upon seed treatment of CAP at 25 μM, the two At*CcSHMT1*-OE lines germinated normally, while the WT plants were severely injured and did not survive beyond 7 days post-treatment (Supplemental Fig. 8S). In line with the increased expression of *SHMT1* in At*CcSHMT1*-OE lines, SHMT concentrations were also elevated in At*CcSHMT1*-OE plants compared to WT plants after CAP treatment, although the concentration of SHMT decreased when high concentrations of CAP were applied (Fig. 4C). Interestingly, the phenotype of the Os*CcSHMT1*-OE lines was similar to that of the At*CcSHMT1*-OE lines under CAP treatment. Upon treatment with CAP at 2.78 mM, the two independent Os*CcSHMT1*-OE lines did not show visible injury, whereas the WT plants were severely injured and did not survive beyond 7 days post-treatment (Fig. 4D). For the two independent Os*CcSHMT1*-OE lines, the EC_50_ values of CAP were 2.415 mM (Os*Cc*SHMT1*-* OE20) and 2.766 mM (Os*Cc*SHMT1*-*OE26), whereas that EC_50_ of WT plants was 1.675 mM (Fig. 4E, Supplemental Table 4S). With the overexpression of *SHMT1* in Os*CcSHMT1*-OE lines, SHMT activity was also elevated in Os*CcSHMT1*-OE plants compared to that in WT plants after CAP treatment, although the activity of SHMT was similar between *Os*SHMT1-OE and WT plants without CAP treatment (Fig. 4F). Taken together, our results indicate that *CcSHMT1* overexpression enhances CAP-resistance in both *Arabidopsis* and rice, further supporting the hypothesis that *Cc*SHMT1 may be the binding target of CAP.

**Fig. 6.**
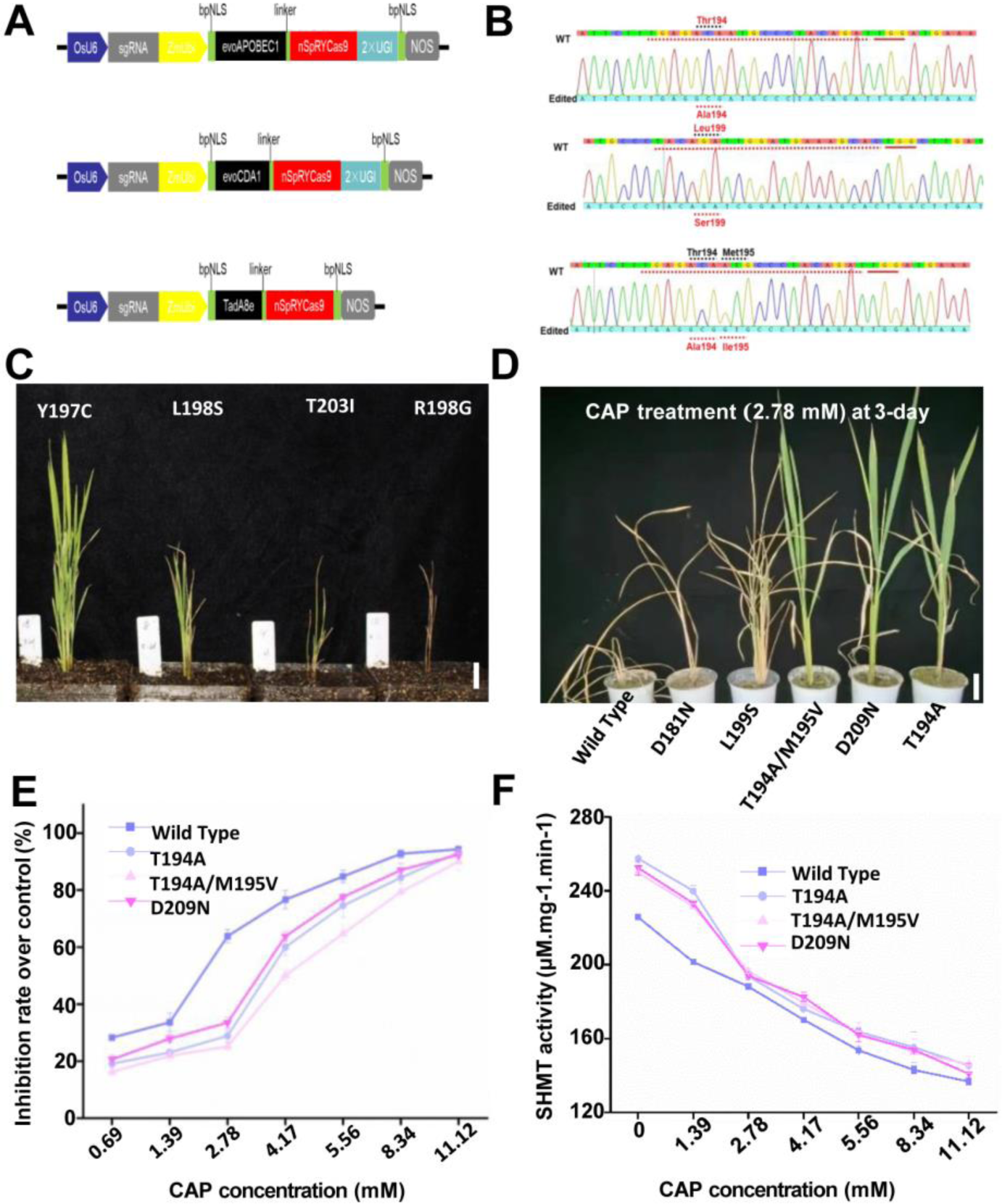
Target evolution of *OsSHMT1* for CAP-resistant mutations by base editor. (A) Architectures of evoCDA1-BE4max, evoAPOBEC1-BE4max, ABE8e, and ACEs. Linker, a 32-aa linker; bpNLS, bipartite nuclear localization signal; TadA8e contains a V106W amino acid substitution. (B) Sanger sequencing chromatograms of four resistant mutants evolved by ACE5. Lower sequences are corresponding target sgRNAs for the mutants. The PAM motif is marked in bold and the boxes indicate the mutated sequences corresponding to the amino acid changes. (C) The typical phenotype *OsSHMT1* mutant lines (Y197C, L198S, T203I, R198G). Bar= 2cm (D)The five lines of mutant plants (D181N, L199S, T194A, T194A/M195V and D209N) and WT 3 days after 2.78 mM CAP treatment. Bar= 5cm (E) The inhibition rate of rice mutant and WT seedlings treated with CAP (0.69, 1.39, 2.78, 4.17, 8.34 and 11.12 mM). The inhibition rate was calculated after 3 days relative to the untreated control plants. (F)The SHMT activity of *CcSHMT1*-overexpressing and WT seedlings treated with CAP (2.78 mM). (G) The influence of CAP (2.78 mM) on SHMT1 content in rice mutants and the wild type by western bolt. Data are the mean±standard deviation (s.d.). Statistically significance was at *p*-value < 0.05 according to Student’s t-test, *P<0.05, **P<0.01, ***P<0.001 and NS, not significant (P>0.05).

### Overexpression of *AtSHMT1* in *Arabidopsis* and *OsSHMT1* in rice boosted resistance to CAP

As previously reported, the photorespiratory *shmt* mutants grow slower than WT plants under normal atmospheric CO_2_ conditions (Moreno et al. 2005). To further confirm SHMT1 is the binding target of CAP, we cloned *AtSHMT1* and overexpressed it in *Arabidopsis*. RT-qPCR assays showed that the transcription levels of *SHTM1* in the two *At*SHMT1-OE lines (*AtSHMT1-*OE7 and *AtSHMT1-*OE14) were significantly increased (Supplemental Fig. 9S). Upon treatment with CAP at 115.62 μM, two *At*SHMT1-OE lines did not show visible injury, while the WT plants were severely injured and did not survive beyond 7 days post-treatment (Fig. 5A). For the two independent *At*SHMT1-OE lines, the EC_50_ of CAP was 369.9 μM (*At*SHMT1-OE7) and 329.0 μM (*At*SHMT1-OE14), whereas the EC_50_ for WT plants was 190.8 μM (Fig. 5B, Supple mental Table 3S).

Interestingly, the knock-out of *OsSHMT1* in rice led to mutant lines grow abnormally and show a typical photorespiratory phenotype (Fig. 5C, Supplemental Fig. 10S), similar to the findings of Wang (2015). To further confirm that SHMT1 was the binding target of CAP, we cloned *OsSHMT1* and overexpressed it in rice. RT-qPCR assays showed that the transcription level of *SHTM1* was highly increased in *OsSHMT1-*OE lines (Os*SHMT1-*OE9 and Os*SHMT1-*OE13) (Supplemental Fig. 9S). The *Os*SHMT1-OE had similarly trends with At*SHMT1*-OE under CAP treated. Under CAP treatment, the *OsSHMT1-*OE9 and *OsSHMT1-* OE13 lines were significantly different from the WT (Fig. 5D). Upon treatment with CAP at 2.78 mM, the two independent *OsSHMT1*-OE lines did not show visible injury, whereas the WT plants were severely injured and did not survive beyond three days post-treatment (Fig. 5D). For the two independent *Os*SHMT1-OE lines, the EC_50_ values for CAP were 2.528 mM (*OsSHMT1-*OE9) and 2.783 mM (*OsSHMT1-*OE13), whereas the EC_50_ for WT plants was 1.675 mM (Fig. 5E, Supplemental Table 4S). The SHMT activity in *CcSHMT1*-overexpressing lines also decreased slightly compared to that in WT plants after CAP treatment, although the activity of SHMT was similar between *Os*SHMT1-OE and WT plants without CAP treatment (Fig. 5F). Taken together, our results indicated that *SHMT1* overexpression enhances CAP-resistance in *Arabidopsis* and rice, suggesting that SHMT1 may be the binding target of CAP.

### The T194A, T194A/M195V and D209N mutation in rice *OsSHMT1* all enhanced resistance to CAP

To comprehensively explore whether SHMT1 is the binding target of CAP and to obtain CAP-tolerant rice plants, the *OsSHMT1* was delimited as the target region for mutagenesis by base editing. The 7^th^ extron of *OsSHMT1* was high conservative among the SHMT1 in the other plants (Fig. 2A; Lakhssassi et al. 2019). A total of 29 guides harboring the NGN PAM were designed in 7^th^ extron using CRISPR-GE (Xie et al. 2017) and cloned into evoCDA1, evoAPOBEC1, and TadA8e base editors in the nSpRYCas9 backbone (Fig. 6A, Wang et al. 2022). These base-editing pools generated dozens of transgenic plantlets with 25 different genotypes (Fig. 6B, Supplemental Fig. 11S). Some mutant lines (Y197C, L198S, T203I, and R198G) did not grow normally and died after transplanting into the soil, indicating the lethal effect of these mutations (Fig. 6C). T-DNA free and homozygous mutant lines with the remaining mutations in the T_1_ generation were further evaluated for CAP tolerance and agronomic traits (Supplemental Fig. 12S). Most tested genotypes were not significantly different from the WT (Supplemental Fig. 13S). Only three genotypes (T194A, T194A/M195V, and D209N) exhibited tolerance to CAP, whereas WT plants died 3 days after 2.78 mM CAP treatment (Fig. 6D). For the three independent T194A, T194A/M195V, and D209N mutants, the EC_50_ of CAP was 2.791, 3.378, and 2.514 mM, respectively, whereas the EC_50_ for WT plants was 1.675 mM (Fig. 6E, Supplemental Table 4S). Field agronomic trait test exhibited that the CAP-tolerance of the genotypes did not affect plant height, tiller number, or 1000-grain weight. However, the seed-setting rate decreased significantly, indicating a potential fitness cost for CAP-resistant (Supplemental Fig. 14S). The SHMT activity was also elevated in three independent T194A, T194A/M195V, and D209N mutant plants compared to the WT plants after CAP treatment, although the activity of SHMT decreased when high concentrations of CAP were applied (Fig. 6F). Altogether, *OsSHMT1* T194A, T194A/M195V, and D209N mutations enhanced CAP-resistance in rice, further supporting the idea that SHMT1 may be the binding target of CAP.

### CAP bound to *Cc*SHMT1

To understand how CAP inhibited enzymatic activity of *Cc*SHMT1, docking analysis was employed to reveal favorable binding affinity between CAP and homology modeling structure of *Cc*SHMT1(affinity: Q=-4.52 kcal/mol, rmsd=1.448<3.5) (Fig. 7A, Supplemental Fig. 15S). The hydroxyl-hydrogen atom of Ser190 forms hydrogen bonds with the carboxyl group of CAP at a distance of 2.9 Å. Ala191 and Val192 also formed hydrogen bonds with carboxyl group of CAP, with distances of 2.5 and 2.4 Å, respectively. The short distances between CAP and the homology modeled structure of *Cc*SHMT1 may be responsible for its favorable binding affinity. Interestingly, the Ser190, Ala191, and Val192 were located in the 7^th^ extron of *OsSHMT1*. To further verify the CAP binding sites, the *Cc*SHMT1 mutants by site-directed mutagenesis were expressed in *E. coli* (Supplemental Fig. 16S). Microscale thermophoresis (MST) analysis of the wild type and *Cc*SHMT1 mutants was used to validate the docking results (Fig. 7B). Non-mutated *Cc*SHMT1 and CAP showed high binding affinities (K_d_=12.3 μM). No binding of VA to *Cc*SHMT1 up to 50 μM was detected (Supplemental Fig. 17S). When Ser190 and Val192 were mutated to Ala190 and Leu192, CAP binding was slightly impeded (K_d_= 15.5 μM and K_d_ = 28.00 μM, respectively). In contrast, mutation of Ala191 to Thr191 resulted in a reduced binding (K_d_= 51.9 μM). Furthermore, the *in vitro* enzymatic activity assay results of the mutated enzyme agreed with those of the MST analysis (Fig. 7C). There was no significant difference between Ser190Ala (190.13 μmol phenylserine·min^-1^·mg^-1^ protein) and Val192Leu (197.83 μmol phenylserine·min^-1^·mg^-1^ protein) site mutations on SHMT enzymatic activity, but the Ala191Thr (154.97 μmol phenylserine·min^-1^·mg^-1^ protein) and three-site mutant (166.30 μmol phenylserine·min^-1^·mg^-1^ protein) were decreased when compare with non-mutation *Cc*SHMT1 (193.35 μmol phenylserine·min^-1^·mg^-1^ protein) without CAP treatment. In addition, there was no significant difference between the Ser190Ala and Val192Leu site mutations, but the Ala191Thr site mutation prevented CAP-induced inhibition of *Cc*SHMT1. Collectively, the data indicate that CAP binds to the key site of *Cc*SHMT1 and that Ala191 provides a critical interaction that is disrupted when mutated to Thr191.

**Fig. 7.**
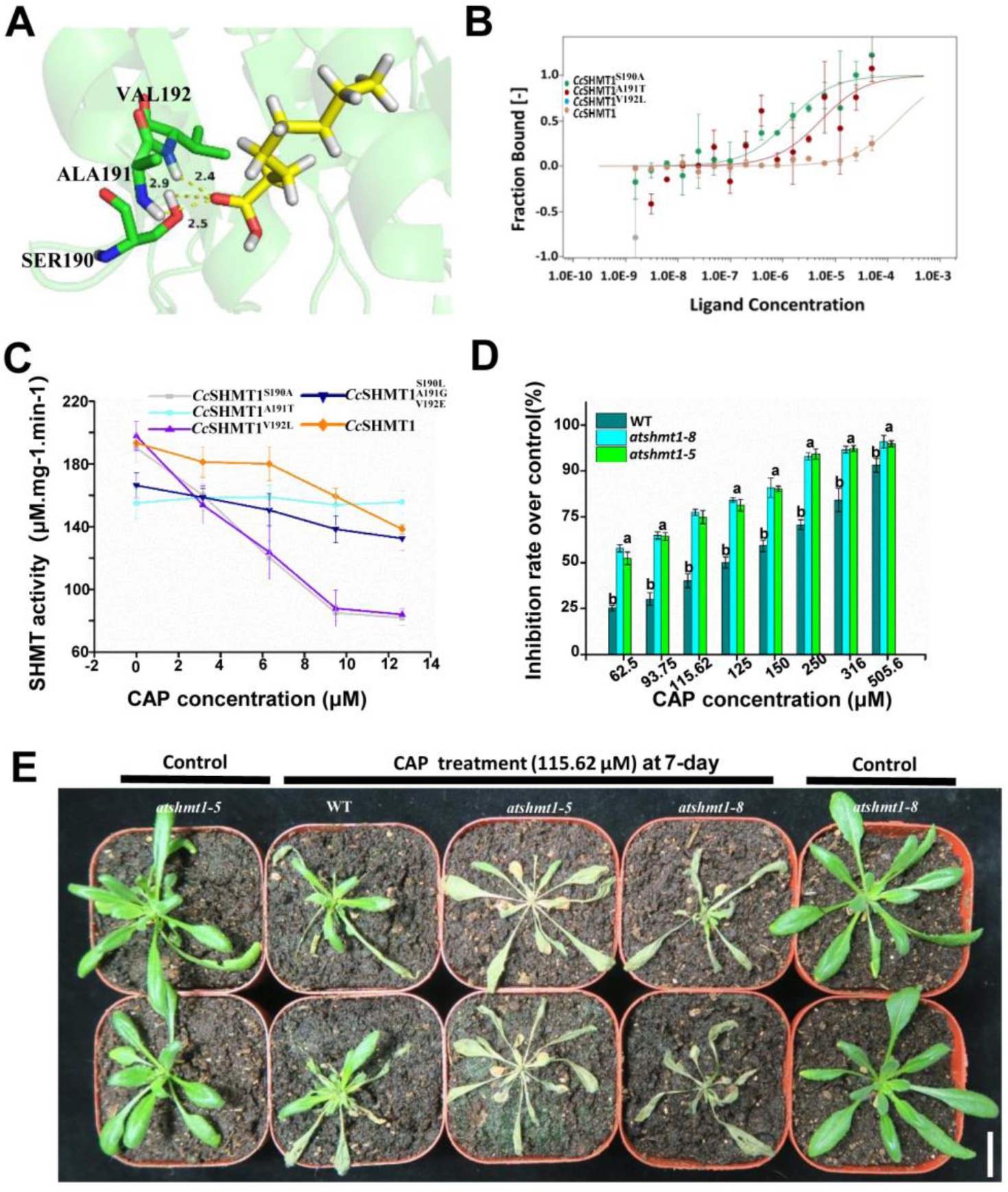
Interaction between *Cc*SHMT1 and CAP. The docking results revealed hydrogen bonding between CAP and homology modeling structure of *Cc*SHMT1 amino acid residues Ala191, Ser190, and Val192. The key residues surrounding the active site are shown as green sticks, and CAP is shown in yellow. Hydrogen atoms of Ser190 were within hydrogen bonding distance of the carboxyl group of CAP (2.5 Å). The hydrogen atom of Ala191 exhibited hydrogen bonds with the carboxyl group at a distance of 2.9 Å. The hydrogen atom of Val192 also exhibited hydrogen bonds with the carboxyl group at a distance of 2.4 Å. The results were produced with PYMOL (V.1.7.0, DeLano Scientific LLC). (B) MST titration profile of the interaction between CAP and *Cc*SHMT1 amino acids. The error bars represent the standard deviation (s.d.) of each data point calculated from three independent thermophoresis measurements. (C) Effect of mutations to *Cc*SHMT1 activity under CAP. *Cc*SHMT1^S190A^: Ser190 was mutated to Ala; *Cc*SHMT^A191T^: Ala191 was mutated to Thr; *Cc*SHMT^V192L^: Val192 were mutated to Leu; *Cc*SHMT1 ^S190L/A191E/V192G^: Ser190, Ala191 and Val192 was mutant to Leu, Gly, and Glu. SHMT activity was assayed for purified recombinant *Cc*SHMT1 proteins by the product rate of benzaldehyde 0.5 h after CAP treatment (3.16, 6.32, 9.48, 12.64 μM). (D) The inhibition rate of two lines of site-directed mutagenesis lines (*atshmt1*-5 and −8), and WT seedlings treated with CAP (62.5, 93.75, 115.62, 125, 150, 250, 316, and 505.6 μM). The inhibition rate was calculated after 7 days relative to the untreated control plants (Inhibition rate % = (Freash weight of untreated control plants-Freash Weight of treatment plants)/Freash Weight of untreated control plants * 100%). (E) The two lines of site-directed mutagenesis lines (*atshmt1*-5, −8, Ser190, Ala191, and Val192 were mutated to Leu, Gly, and Glu) and the WT 7 days after 115.62 μM CAP treatment. Bar=1 cm. The MST analysis and activity assay comprised three biological replicates, each with three technical replicates. Data are reported as mean ± s.d. Results with a *p*-value < 0.05 according to Student’s t-test were considered statistically significant.

Furthermore, we obtained the *At*SHMT1 site-mutants that were site-directed *in vivo*. DNA sequence data verified that the Ser190, Ala191 and Val192 were mutated to Leu, Gly, and Glu in the independent site-directed mutagenesis lines (−5 and −8) (Supplemental Fig. 18S). To determine the optimal concentration of CAP for the treatment of *Arabidopsis*, site-directed mutants and WT seedlings were treated with CAP at different concentrations. Upon treatment with CAP at 62 μM-505μM, the site-directed mutagenesis plants were severely injured than WT beyond seven days post-treatment (Fig.7D). Whereas the site-directed mutagenesis plants were more severely injured than WT beyond seven days post CAP treatment at 115.62 μM (Fig. 7E). Collectively, our results indicated that CAP bound to the target site of *Cc*SHMT1 and that Ser190, Ala191, and Val192 are critical amino acid residues for *Cc*SHMT1 interaction with CAP.

## DISCUSSION

Uncovering the binding targets of CAP is a crucial step in developing commercial herbicide and herbicide-resistant crop. Combinatorial analysis of untargeted metabolomic and proteomic profiles revealed nine candidate proteins involved in eight different pathways. Among these candidate proteins, SHMT1 had the highest frequency in the top eight KEGG pathways (Fig.1C). *Cc*SHMT1 also showed the highest significantly changed in CAP-responsive expression (Fig. 3A). Several lines of evidence suggest that *Cc*SHMT1 serves as a target for CAP herbicidal activity in *C. canadensis*. First, overexpression of *Cc*SHMT1 conferred CAP resistance in transgenic *Arabidopsis* and rice, suggesting that excess *Cc*SHMT1 is the direct target of CAP (Fig. 4). Second, both *in vivo* and *in vitro* experiments confirmed that CAP inhibits SHMT enzyme activity, and this inhibition is competitive, as shown by *in vitro* kinetics assays (Fig. 3D-F). Third, the mutant plants harboring T194A, T194A/M195V, and D199N mutations exhibited tolerance to CAP, whereas the WT plants died after CAP treatment (Fig. 6C). Four, structural modeling suggested high affinity binding of *Cc*SHMT1 to CAP, and the critical amino acids for this interaction were experimentally verified (Fig. 7). Our experimental evidence suggests that *Cc*SHMT1 is an authentic binding target of CAP, which explains its herbicidal activity against *C. canadensis*.

The amino acid biosynthesis enzymes are one of the most promising herbicide target (Heap 2023). SHMT1, which plays a role in photorespiration and metabolic conversion of glycine to serine, is also an important enzyme for survival or death in response to biotic and abiotic stresses in plants (Hu et al. 2022; Liu et al. 2019; McClung et al 2000). In *Arabidopsis*, *AtSHMT1* (At4g37930) is necessary and sufficient for SHMT-dependent photorespiration and controls abiotic stress-induced cell damage (Voll et al. 2006, Moreno et al. 2005). Our data showed that knock out or base editing of *OsSHMT1* in rice led to the abnormally growth of mutant lines and a typical photorespiratory phenotype (Fig. 5C). Similarly, photorespiratory *shmt* mutants grow slower than WT plants under normal atmospheric CO_2_ conditions and are hypersensitive to abiotic stress (Moreno J.I. et al. 2005). Meanwhile, CAP is a competitive inhibitor of *Cc*SHMT1 with Ser (Fig.3), and has a structure similar to that of CAP and Ser, resulting in leaf chlorosis and plant death. Indeed, a slow inhibition rate could herald comparatively high dosages. The recommended usage of CAP in the field was further confirmed by the SHMT activity inhibition rate data (Fig.3E). Similarly, the comparatively lower dosage of branched-chain amino acid biosynthesis inhibitors were significantly specific inhibited the binding targets, such as acetolactate synthase (ALS, EC 2.2.1.6) (Dezfulian et al. 2017; Gil-Monreal et al. 2019; Fuchs et al. 2021). Therefore, SHMT1 is an amino acid biosynthesis target and get inhibited by CAP.

Similarly, determination of the binding site is essential for the discovery of novel herbicides target and the creation of herbicide-resistant crops (Pfister and Arntzen 1979; Schloss 2010). Molecular docking predicted how CAP fits into the binding site of homology modeling structure of *Cc*SHMT1, and the amino acid Ala191 was found to be essential for CAP binding (Fig.7). Surprisingly, Ala191 is located at N-terminal domain in the *At*SHMT2 structure (Nogues I et al. 2022). Similarly, the Thr183, Asn356, and Thr357 residues in Plasmodial SHMT, also located in the N-terminal domain, establish strong hydrogen bonds with the pyran ring of carboxylate (2-(1-(3-(6-amino-5-cyano-3-methyl-2,4-dihydropy rano[2,3-c]pyrazol-4-yl)-7-fluoro-2,2-dimethyl-2,3-dihydro-1H-inden-5-yl) piperidin-4-yl)acetic acid). Such alternative bonds to the pyran ring can potentially be introduced in novel SHMT inhibitors (Schwertz G et al. 2018). More recently, the AS-IE method was applied to analyze the binding pocket of SHMT2 to 28 different SHMT2-inhibitor complexes. Tyr105 and Arg425 are also located in the N-terminal domain and have been found to be major contributors to the total binding free energy (He LP et al. 2019). These previous reports show that the binding sites of CAP are located at the key domains of known SHMT monomers, that is, the Ser-binding pocket at the N-terminus. Interestingly, three genotypes (T194A, T194A/M195V, and D209N) exhibited tolerance to CAP while WT plants died three days after 2.78 mM CAP treatment (Fig. 6D). The T194 and D209 are also located in the 7^th^ extron of the N-terminal domain. Hence, the 7^th^ extron of SHMT1 at N-terminal domain is the key binding site of CAP, which reveals a novel mode of action and herbicide-resistant crop.

Multiple separate targets are widespread in the natural compounds (Chen HS et al. 2017). One hypothesis might explain how multiple changes occur in weeds and the low level of resistance in crops, which is possible because CAP has multiple separate targets in weeds. Unexpectedly, two *Cc*SHMT1 overexpressing lines enhanced the resistance level, whereas the site-directed mutagenesis plants were more severely injured than the WT plants seven days post-treatment with CAP (Fig. 5G). Normally, site-directed mutagenesis of the active site confers tolerance to the inhibitor as the binding affinity decreases (Kandoth PK et al. 2017). However, some studies have reported that mutations lead to sensitivity (Beard and Wilson 1994). Similarly, the signaling-sensitive M637W, S641A, and Y667A thyroid-stimulating hormone receptor (TSHR) exhibited 25-50% activity compared to the WT and led to a constitutively active receptors (Kleinau G et al. 2010). *A. thaliana shm1-1* mutants are more susceptible to infection by biotrophic and necrotrophic plant pathogens and accumulate more H_2_O_2_ than WT plants under salt stress (Moreno et al., 2005). Indeed, the structures of all known SHMTs are extensively similar, and are either homodimers or homotetramers of monomers with similar molecular masses (Nogues I et al. 2022). Obviously, the functional redundancy of SHMT is commonly observed in plants under stress (Anderson DD et al. 2009). The 3D model of SHMT could be docked with various substrates, inhibitors and co-factors alone or in combination to analyze the structural change(s) in different SHMT complexes (Trivedi et al. 2002). The central role of SHMT isozymes in producing one-carbon-substituted folate cofactors suggests that these enzymes (isozymes) influence photorespiration and therefore could serve as herbicide targets (Witschel 2012). SHMT1 is a member of the SHMT family, which has five SHMTs in rice and seven SHMTs in *Arabidopsis*; thus, there might be redundant proteins of SHMT1 (Moreno et al. 2005; Yan MY et al. 2022). Therefore, the *Cc*SHMT1 is the binding target of CAP and the other targets of CAP may exist.

Most successful long-established targets involve more than one molecule or possess targets that are used against multi-resistant weeds (Gray and Wenzel 2020). In the case of *C. canadensis*, multiple resistance has been reported, making it difficult to control this weed (Heap, 2023). For instance, mutations in the 5-enolpyruvylshikimate-3-phosphate synthase (EPSPS) gene, such as P106T, and mutations in the acetolactate synthase (ALS) gene, such as P197A, have been identified (Amaro-Blanco I et al., 2018; Mora DA et al., 2019). These studies on target sites are very well done to identify herbicide targeting amino acid sequences in natural populations. In our study, we obtained mutant populations by base-editing the 7^th^ exon of the *Os*SHMT1 gene. Fortunately, three genotypes (T194A, T194A/M195V, and D209N) exhibited tolerance to the herbicide CAP, while wild-type plants died three days after treatment with 2.78 mM CAP (Fig. 6D). Although more variations of SHMT1 are needed to determine whether they confer higher CAP-tolerance, this target site mutant also supports the herbicide targeting mechanism effectively. Similarly, mutations in the EPSPS gene, such as D213N, and mutations in the ALS gene, such as P171F, confer reliable resistance to glyphosate and nicosulfuron in rice (Zhang R. et al., 2021; Zhang C. et al., 2023).

Interestingly, SHMT1 is involved in six out of the eight pathways in *C. canadensis*, including carbon fixation in photosynthetic organisms, amino acid biosynthesis, and carbon metabolism (P<0.05) (Fig. 1C and 1D). The targets it possesses are encoded by multiple genes (Brazel and Maoileidiah 2019). Compounds that bind to multiple proteins as targeting mechanisms make the evolution of target-site resistance mechanisms more challenging (Gaines TA et al., 2020). Therefore, SHMT1 can be utilized to combat multi-resistant weeds and is a promising target for herbicide development and herbicide-tolerant crops.

In summary, CAP competitively binds to the key domain of *Cc*SHMT1, disrupting SHMT function and resulting in the inhibition of *C. canadensis* growth (Fig.8). The studies presented herein provide a novel integral framework for herbicide binding target identification and the development of novel SHMT-targeting inhibitors. In the future, CAP-based inhibitors that can inhibit all SHMT isoforms or other target proteins as well as CAP-tolerant crops will be developed.

**Fig. 8.**
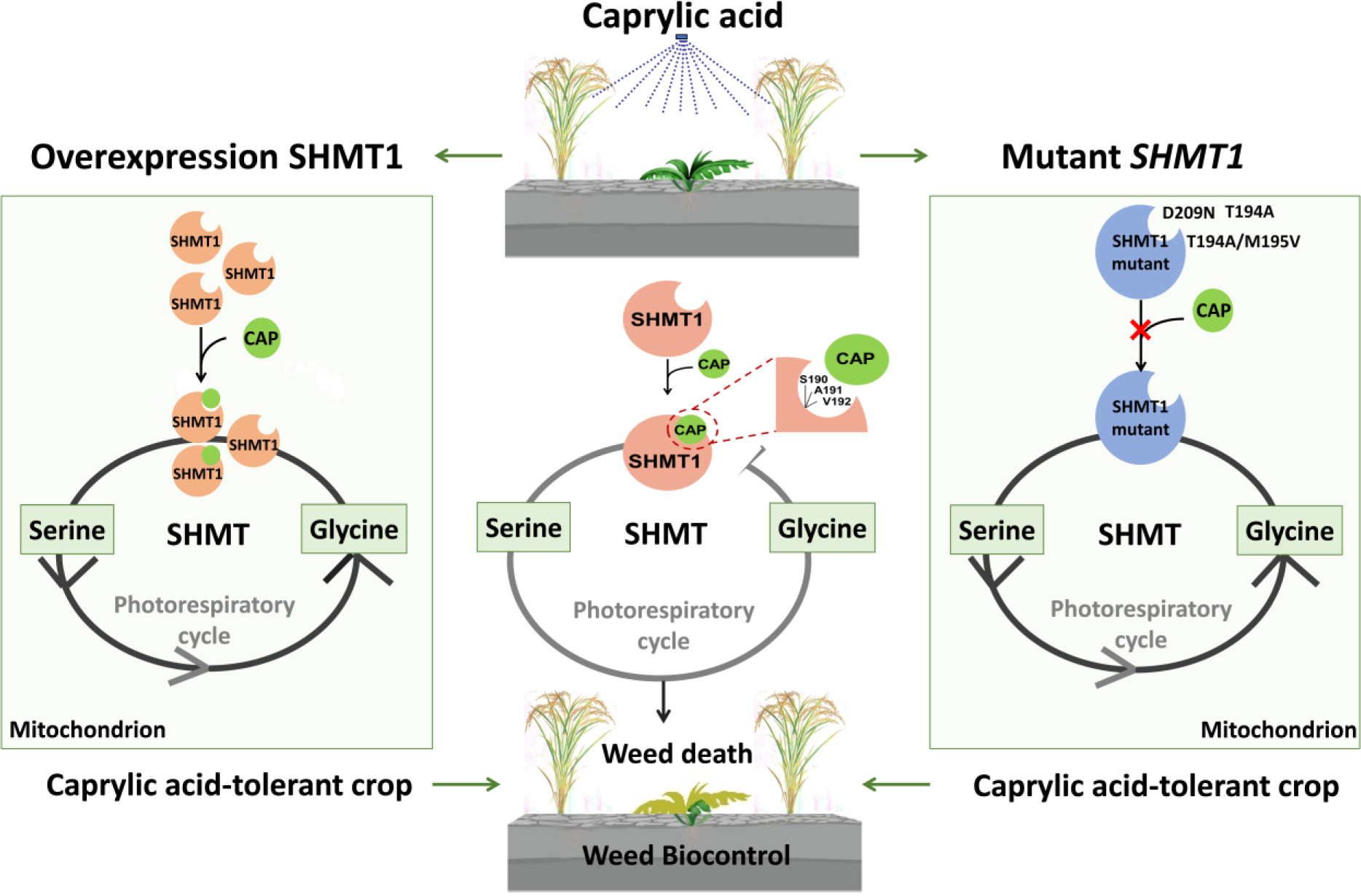
Model of Caprylic acid inhibition of SHMT1 to subdue weeds and tolerant crop. Under normal conditions, SHMT1 catalyzes the reversible interconversion of serine and glycine. The non-mutation *Cc*SHMT1 showed good binding affinities with caprylic acid. In normal *CcSHMT1* plants, caprylic acid competitively binds to the key domains (Ser190, Ala191, and Vla192) of *Cc*SHMT1, thus disturbing SHMT function and resulting in the inhibition of *C. canadensis* growth. In the overexpression *CcSHMT1* plants, caprylic acid competitively binds to the key site of *Cc*SHMT1. However, serine can still be converted to glycine in surplus *Cc*SHMT1, resulting in higher crop survival rates. In the mutant *OsSHMT1* plants, we obtained the T194A, T194A/ M195V, and D209N site mutant lines. Caprylic acid cannot competitively bind to the key site (Ser190, Ala191, and Vla192) of *Cc*SHMT1. Serine can be converted into glycine, resulting in higher crop survival rates. SHMT1: Serine hydroxymethyltransferase 1.

## Materials and Methods

### Plant material

*C. canadensis* seeds were collected from a recreational area in Changsha, China (N28°28′21″, E113°4′52″). *C. canadensis* and rice plants were grown in an artificial climate chamber with a 16 h photoperiod (100-120 μmol m^-2^ s^-1^) and 22/18 °C (day/night temperatures). *A. thaliana* plants were grown in a chamber with a 16 h photoperiod (100-120 μmol m^-2^ s^-1^) 23/20 °C (day/night temperatures) and 65% relative humidity. *Arabidopsis* accession Col-0 was used as the WT. Plants at the stage of 3 to 5 leaves were used in the experiments. The primers and methods used for mutant genotyping are listed in Supplemental Table 5S.

### Leaf untargeted metabolomics based on GC-MS

*C. canadensis* were treated with 625 μM CAP using a spraying tower. At 4 h after CAP treatment and untreatment, a total of 2 g of leaves (8 technical replicates) were collected for analysis. Each leaf sample was analyzed using an Agilent 7890B gas chromatograph coupled to an Agilent 5977A mass-selective detector (MSD) (Agilent Technologies Inc., CA, USA). Metabolites were unambiguously identified using the National Institute of Standards and Technology (NIST) and Fiehn Lab databases and quantified according to the peak height of a unique ion. The data were subjected to PCA and (orthogonal) partial least squares discriminant analysis (OPLS-DA). Differentially expressed metabolites were selected based on a combination of VIP values larger than one and *p*-values less than 0.05. Additional details regarding preprocessing, GC-MS analysis, statistical analysis and identification of differential metabolites are shown in Supplemental Table 2S.

### Combinatorial pathway analysis of proteomics and metabolomics data

The proteomics and metabolomics data at the 4h time point were chosen for the analysis of common pathways and identification of potential protein targets of CAP in *C. canadensis*. The metabolite IDs were transformed into the KEGG ID format using MBROLE (http://csbg.cnb.csic.es/mbrole2/). Additional details of the KEGG pathway and Gene Ontology (GO) analyses are provided in the Supplemental Marterial.

### SHMT *in vivo* enzyme assay and qPCR analysis

The relative expression levels of the *CcSHMT1* gene were analyzed by RT-qPCR 30 min after CAP treatment (625 μM). Additional different genes including *FBA5*, *PGK3*, *At1g56190*, *PGM1* and *GLO1* were tested using the method described for *SHMT1*. The detail of qPCR are shown in the Supplemental marterial.

At 0, 0.5, 1, 2, 4, and 8 h after 625 μM CAP treatment, *C. canadensis* leaf materials (2 g) were collected for analysis of the relative expression of *CcSHMT1* by qPCR and *Cc*SHMT1 protein levels with a SHMT ELISA Kit (Mei5 Biotech., Beijing). Additionally, at 4h after treatment with 0, 78.125, 156.25, 312.5, 625, or 1250 μM CAP, *C. canadensis* leaf materials (2 g) were collected for SHMT activity measurement as previously described (Waditee-Sirisattha R et al. 2012). Each independent measurement was consisted of at least two replicates.

### Cloning and bioinformatics analysis of *C. canadensis SHMT1*

The *SHMT1* CDS was cloned directly from *C. canadensis* mRNA by RT-PCR using primer-C (Supplemental Table 5S). To obtain the full length cDNA of *CcSHMT1*, 5′ Rapid amplification of cDNA ends (RACE) was performed using the primer-R5 (Supplemental Table 5S) with the 5′ Full Race Core kit (TAKARA, Japan). The bioinformatics analysis of *CcSHMT1* sequences show in Supplemental marterial. The homology modeling structure of *Cc*SHMT1 proteins was performed using AlphaFold2 by RoseTTAFold (Baek MY et al. 2021). PyMOL molecular viewer was used to generate the output before selecting a rational conformation.

### Subcellular Localization of *CcSHMT1*

To construct the 16183-*CcSHMT1*-GFP vector, the total CDS of *CcSHMT1* gene sequence (1539-bp) was obtained as described above and cloned into the 16318-GFP. Mesophyll protoplasts were prepared from the leaves of 4-week-old *Arabidopsis* and transformed by the polyethylene glycol method (Goyal and Katiyar 1994). The subcellular localization was co-localizated with GFP fluorescence overlapping with Mito fluorescence (M7512, Thermo Fisher Scientific), demonstrating the mitochondrial localization of SHMT. The subcellular distribution of *CcSHMT1* was observed using a confocal laser scanning microscope (Leica, Germany).

### Expression and purification of *Cc*SHMT1 protein and *in vitro* enzyme activity assay

The total CDS of *CcSHMT1* gene sequence (1539-bp) was obtained as described above. The PCR-amplified fragments using primer-EX (Supplemental Table 5S) were purified using a kit (Sangon Biotechnology Co., Ltd., China) and ligated into the pET28a vector using ExnaseII (ClonExpress II One Step Cloning Kit, Vezyme Biotechnology Co., Ltd.). The details of the expression and purification of *Cc*SHMT1 protein are shown in the Supplemental marterial. A 20 μL aliquot of *Cc*SHMT1 protein in PBS buffer at various concentrations (0.075-0.117 mg/mL) were used in the assays. The SHMT enzyme activity was measured as previously described (Waditee-Sirisattha R et al. 2012). To measure the kinetic parameters (K_M_ and V_Max_) of SHMT with CAP or VA (18.9 μM), different dosages of DL-β-phenylserine (1 mM, 1.25 mM, 2.00 mM, 2.5 mM, 5 mM, 10 mM) were used. The kinetic parameters were calculated using the Michaelis-Menten equation (Nahler 2004). The SHMT reaction was assayed 30 min after CAP treatment. Each independent measurement consisted of at least two other replicates.

### Overexpression of *CcSHMT1* or *AtSHMT1* in *Arabidopsis* and rice

The pCambia2301-CcSHMT1 or pCambia2301-AtSHMT1 plasmids were then transformed into *Agrobacterium tumefaciens* GV3101 or EH105 strains. *Arabidopsis* plants were transformed using the floral dip method. The *Agrobacterium*-mediated transformation was performed as previously described (Wang MG et al., 2019). The homozygous T2 lines were selected for further analysis.

Transgenic *Arabidopsis* seeds were grown on agar plates (40 mL) containing CAP at five concentrations (0, 3.16, 6.32, 15.8, 25 and 31.6 μM) to test CAP sensitivity during seed germination. The five-leaf stage of homozygous T2 *Arabidopsis* was sprayed with diluted CAP at five concentrations (0, 189.6, 316, 505.6, 632 and 1264 μM) to test CAP sensitivity. The five leaves stage of homozygous T2 *Nipponbare* were sprayed with diluted CAP at five concentrations (0.69, 1.39, 2.78, 4.17, 8.34 and 11.12 mM) to test CAP sensitivity. The bioactivity assay comprised three biological replicates, each containing three technical replicates. The *SHMT1* gene expression and enzyme activity was performed as described above.

### Evolving *OsSHMT1* for CAP-tolerance mutant

Target sequences within the 7^th^ exon of *OsSHMT1* were subjected to CRISPR-GE (Xie et al., 2017) for guide generation. The guide sequences were scanned for the presence of NGG or NRN (R=G/A) PAM motifs on both the sense and anti-sense strands. In total, 29 guides were selected for assembling the sgRNA library. The oligos of the guides were synthesized by Sangon Biotech (Shanghai, China) and cloned into the evoCDA1, evoAPOBEC1 and TadA8e (V106W) base editors as described by Wang et al. (2022). All primer sets used are listed in Supplementary Table 5S. The base editing plasmid library was introduced into *A. tumefaciens* strain EHA105 and transformed into embryogenic calli of Xiushui134 (*Oryza sativa* L. Japonica). Transgenic plants were screened using 40 mg/mL hygromycin, and the positive plants were transferred to the soil. Homozygous and T-DNA free edited lines were identified in the T1 generation and further propagated into the T2 generation. The homozygous and T-DNA free transgenic lines and wild type lines were treated by the dosage of CAP (0.69, 1.39, 2.78, 4.17 and 8.34 mM) under the greenhouse conditions. After 3 days of treatment, the aboveground parts of the plants were calculated. The bioactivity assay comprised three biological replicates, each with three technical replicates.

### Binding affinity measurement

To illuminate structural and functional relevance between CAP and *Cc*SHMT1 protein, docking studies were carried out with AutoDocking Tools-1.5.6 according to a previously described methodology (Forli et al. 2016). This binding model was chosen for further experiment, since it had the lowest binding energy and an RMSD of <3.5. Three destabilizing mutations (putative active-site residues, Ser190, Ala191, and Val192) were introduced by mutagenesis PCR using primer-M (Supplemental Table 5S). MST was recorded using the MO control software, and the binding affinities were analyzed using PALMIST and GUSSI (Chad Brautigam, UT Southwestern, Dallas, TX). The mutant protein enzyme activity *in vitro* assay was performed according to the above description.

### Site-directed mutagenesis and transformation

The *At*SHMT1 (AT4G37930) amino acid substitutions were generated using a site-directed mutagenesis kit protocol (Stratagene). The primer-SD was used for Ser190 Leu, Ala191 Gly and Val192 Glu in supplemental Table 5S. The pCambia2301-KY-SHMT1 plasmid was then transformed into *A. tumefaciens* GV3101 strain, and *Arabidopsis* plants were transformed by the floral dip method. Homozygous T2 lines were selected for further analysis. Homozygous T2 plants were sprayed with diluted CAP at five concentrations (0, 62.5, 93.75, 115.62, 125, 150, 250, 316, and 505.6 μM) to test CAP sensitivity during plant growth. The bioactivity assay comprised three biological replicates, each with three technical replicates. *AtSHMT1* expression was analyzed in the transgenic and WT plants using qPCR. The mutate sites of homozygous T2 plants were performed to verify correctness by DNA sequencing.

### Statistical analyses

Data from the representative experiment each are presented. Most data provide the mean values ± standard deviation (s.d.). Statistical analysis was performed using the analysis of variance (ANOVA) test and was considered statistically significant at *p*-value < 0.05, according to the Student’s t-test.

## Acknowledgments

The authors thank Dr. Qing Chang for MST data collection at the Protein Preparation and Characterization Platform of Tsinghua University Technology Center for Protein Research. This work was supported by the Natural Science Foundation of China (32172433), Natural Science Foundation of Hunan Province (2021JJ20034), the China Agriculture Research System of MOF and MARA (CARS-16-E19), the Hunan Agriculture Research System (2022-31), and Scientific-Innovative of Hunan Agricultural Sciences and Technology (2022CX70, 2021CX42 and 2022CX11).

## Author’s contribution

Lianyang Bai and Zuren Li designed the experiments and wrote the article. Zuren Li, Haodong Bai, Hongzhi Wang, Jincai Han, Likun An, Dingfeng Luo, Yingying Wang and Wei Kuang performed experiments. Zuren Li and Mugui Wang analyzed the data. All authors commented on the manuscript.

